# A mathematical sequence representing tonic action potential spike trains

**DOI:** 10.1101/2024.07.23.604868

**Authors:** Dongil Keum, Kwon-Woo Kim, Molly Pruitt, Alexandre E. Medina de Jesus

## Abstract

This is a study outlining the regularity of action potential spikes. Through a stochastic study, we observed a series of strong correlations between the intervals of tonically firing spikes generated by injecting constant currents of varying intensities into layer V pyramidal neurons of the ferret medial prefrontal cortex. Based on this, we derived a formulaic relationship for the interspike intervals (ISIs). According to this formula, an ISI can be expressed as a product of two factors: the timing precursor and the scale factor. Those arise from a linear relationship between activities of ion channels that modulate spike frequency adaptation and spike timing. Using this rule, we successfully predicted spike timing and demonstrated that the spike timing can be determined by the linear combination of various ion channel activities, reflecting different cellular signaling pathways such as G-protein coupled receptor (GPCR) activation. These findings not only aid studies on cellular signaling but also expand our insight into neural coding, while increasing research efficacy through neural modeling.

**Significant Statement:** While the action potential (AP) pattern may appear simple at first glance, no rule has been discovered in the nearly 100 years since it was first recorded. Building on this finding, we have developed a method to intuitively measure the activity of various ion channels responsible for determining spike timing from the AP spikes, as well as the associated intracellular and extracellular signaling pathways.

## Introduction

Spikes are the fundamental unit of communication between neurons. A neuron fires spikes due to its behavior as a resistor-capacitor (*RC*) circuit with time- and voltage-dependent ionic conductance, originating from actions of different voltage-gated ion channels and their corresponding electromotive sources, which are derived from ion gradients between the plasma membrane (Hodgkin and Huxley, 1952). Neurons receive spikes from other neurons as input and fire spikes when those inputs evoke an appropriate voltage change (Geisler and Goldberg, 1966; Roy and Smith, 1969; Abbott, 1999; Lapicque, 2007). As the stimulus becomes stronger, the number of spikes increases. However, the timing of spike occurrence varies dynamically for each spike even with the same stimulus due to many factors such as the stochastic opening and closing of ion channels following the Boltzmann distribution.

The spike timing, along with the firing rate, plays an important role in neural functions. Precise spike timing is crucial for processing sensory input (Bohte et al., 2002). The temporal interval between pre- and post-synaptic spikes is the basis of spike timing-dependent plasticity (STDP), as their order can result in long-term potentiation (LTP) or depression (LTD) (Feldman, 2012; Andrade-Talavera et al., 2023). The spike frequency adaptation (SFA), which is an intrinsic cellular function that delays spike timing when a stimulus persists, is crucial for sensory processing in the lobula giant movement detector (LGMD) of locusts, and is possibly an underlying mechanism of how cholinergic signaling modulates working memory (Gu et al., 2005; Galvin et al., 2020). Furthermore, computational studies suggest that SFA is an important component of working memory formation (Salaj et al., 2021), language processing (Fitz et al., 2020), and spike neural networks (Salaj et al., 2021; Ganguly et al., 2024) where short-term memory and neuronal synchronization are required.

Spiking is assumed to be a random renewal process (Cox and Lewis, 1966) in which each spike is an independent event. However, it is frequently observed that spikes reflect previous spiking history (Quirk and Wilson, 1999; Keat et al., 2001) and their interspike intervals (ISIs) are correlated over several spikes in vivo (Kuffler et al., 1957; Ratnam and Nelson, 2000; Avila-Akerberg and Chacron, 2011; Schwalger and Lindner, 2013; Ramlow and Lindner, 2021). This phenomenon contributes to signal detection (Ratnam and Nelson, 2000; Nesse et al., 2021) and information filtering (Sharafi et al., 2013), and, based on this, studies are conducted to accurately simulate spike occurrence (Keat et al., 2001; Truccolo et al., 2005).

As a first step toward our long-term goal of understanding how G protein-coupled receptor (GPCR) signaling influences neural coding, especially in working memory and multisensory processing, the objective of this study is to identify rules that determine spike patterns. When a constant current was injected, the pyramidal neurons produced tonic spikes. Despite the appearance of a pattern in tonic spiking at first glance, no rules have been discovered thus far. We observed that all ISIs exhibited a strong Pearson’s correlation coefficient with adjacent ISIs. This relationship was described by a formula in which the ISI changes according to the spike order. According to this formula, an ISI is determined by the timing precursor, which is a recursive number sequence that carries the history of the previous spikes, multiplied by a scale factor that originates from the membrane capacitance. This allowed us to decipher the factors that determine spike timing, which can be explained by a biophysical model with Hodgkin-Huxley (HH) like conductance. We observed that the repolarization conductance of the spike increases as the spikes repeat, and this is proposed to be due to the intervention of various ion channels that regulate SFA. This conductance delays the rise in membrane potential by adding to the leak current until the neuron fires the next spike. Additionally, the increased leak current can be expressed as a linear combination of each ion channel factor, potentially enabling the backward trace of the related ion channel activities from the spike.

Thus, the present study is expected to greatly assist in researching the effects of cellular signaling, such as that of lipids, GPCRs, and calcium, on spike timing, and to significantly impact the improvement and treatment of related neurological disorders. Additionally, it can be utilized to circumvent previous complex neuron models, potentially reducing the technical burden and cost of neural signal detection, neural coding, and neural network research.

## Methods

### Animal perfusion and brain slice preparation

All procedures were performed in compliance with the Institutional Animal Care and Use Committee of the University of Maryland. Two types of artificial cerebrospinal fluid (aCSF, modified from Ting et al., 2014) were used in this procedure: (1) the protective aCSF containing (in mM) 92 N-Methyl-D-glucamine (NMDG), 2.5 KCl, 30 NaHCO_3_, 1.25 NaH_2_PO_4_, 20 hydroxyethyl piperazine ethane sulfonic acid (HEPES), 25 D-glucose, 2 thiourea, 3 Na-pyruvate, 3 Na-ascorbate, 0.5 CaCl_2_ and 10 MgCl_2_, and (2) the holding aCSF, containing 92 NaCl, 2.5 KCl, 30 NaHCO_3_, 1.25 NaHPO_4_, 20 HEPES, 25 D-Glucose, 2 thiourea, 3 Na-pyruvate, 5 Na-ascorbate, 2 CaCl_2_ and 2 MgCl_2_. Ferrets (mustela putorius puro, Marshall Farms, North Rose, NY) were euthanized with isoflurane and perfused with a chilled (4°C) protective aCSF for 4 to 6 minutes, until clear buffer appeared from the drain (30-40 ml/min, 200–250 ml). After decapitation, coronal slices (400 μm) containing both prelimbic and infralimbic cortices (PrL and IL, respectively) were sectioned in the chilled protective aCSF with a vibratome (VT1000s, Leica Biosystems) and then recovered in warm protective aCSF (32°C, 5min) before being transferred to holding aCSF. Slices were kept in the holding aCSF for at least 1 hour for recovery and moved to the recording chamber. All aCSFs in this study were oxygenated with 95% O_2_, 5% CO_2_.

### Patch clamp electrophysiology

Slices were placed on the recording chamber and observed with Olympus BX51 upright microscope equipped with an sCMOS camera (Rolera Bolt, QImaging). Whole cell patch clamp recordings were performed from visually identified pyramidal neurons within layer V of the prelimbic cortex (PrL). Recordings were made in current clamp mode and cells with an access resistance (Ra) of >25 MOhm were excluded from the analysis. The recording aCSF contains 126 NaCl, 2.5 KCl, 26 NaHCO_3_, 1.25 Na_2_HPO_4_, 2 MgCl_2_, 2 CaCl_2_, 10 Glucose, 1 thiourea, 1.5 Na-pyruvate and 2.5 Na-ascorbate and perfused at a speed of 4-5 ml/min. Patch pipettes (4–6 MΩ) were pulled from borosilicate glass capillaries using a PC-10 (Narishige) vertical puller. The pipettes were filled with internal pipette solution consisting of the following (in mM): 130 K-gluconate, 4 KCl, 10 HEPES, 0.3 ethylene glycol-bis(β-aminoethyl ether)-N,N,N′,N′-tetra acetic acid (EGTA), 10 phospocreatine-Na_2_, 4 Mg-ATP, and 0.3 Na_2_-GTP, titrated to pH 7.4 with KOH and adjusted to 285-290 mMol/kg with sucrose. Signals were acquired by a MultiClamp 700B amplifier then digitized at 10 kHz and low-pass filtered at 3 kHz with an 8-pole Bessel filter using a Digidata 1550B and Clampex 11 (Molecular Devices). To generate action potential spikes, step currents were injected from −40 to 340 pA with an increment of 20 pA and duration of 500 ms.

### Experimental design and statistical analyses

Datasets were generated from a total of 27 cells from 8 animals and analyzed and displayed using homemade Python and R scripts. Briefly, *.abf files containing current clamp recordings were loaded using a Python library, pyABF (Harden Technologies), and then the timing and shape of each action potential spike were collected and assigned code for identifying the animal, neuron, current intensity, and spike order. Analysis was performed using R, except for the raster plot shown in Figure 5E. The raster plot was obtained by convolving the actual or predicted timing of the spike train with an asymmetric kernel with τ_decay_ = 3ms.

## Results

In this study, we investigated the regularity of the action potential spikes recorded using the whole cell patch clamp method in 27 layer V pyramidal neurons (PN) of medial prefrontal cortex (mPFC) slices from ferrets aged postnatal day (P) 180 (Fig. 1A). Neurons showing phasic action potentials or generating spikes smaller than 30% of the tallest spike were excluded from the analysis. Action potential spike trains were evoked by 500 ms current steps from −40 pA to 340 pA in 20 pA increments (20 steps). Neurons generated simple tonic action potential spikes showing spike frequency adaptation (SFA) with cell-to-cell variation, but no bursts were observed in our dataset. All neurons fired more than two spikes after 120 pA, and responses at 140 pA, 240 pA, and 340 pA, representing low, medium, and high current, respectively, were selected for convenience of comparison. Pyramidal neurons fired more with stronger stimulation as can be seen in Figure 1B, which shows spikes for current injections as a result of different intensities in a neuron. On average, neurons fire 6.19±0.42, 10.89±0.44, and 12.70±0.43 times at 140 pA, 240 pA, and 340 pA, respectively. We assigned identification codes to all neurons, injected currents, and ISIs to enable hierarchical access. By taking advantage of this, we could examine the change in ISI in individual neurons with that of all neurons (Fig. 1C). Briefly, we accumulated ISI for each neuron and assigned a color code to each ISI order. Then, referring to the existing reports that the first ISI (ISI_1_) was somewhat irregular compared to other ISIs (Guan et al., 2015), it was sorted based on the second ISI (ISI_2_). As a result, we confirmed that as the intensity of the current increases, more spikes occur in the limited stimulation time (500 ms). Interestingly, we observed that when the value of the second ISI increases, the cumulative value up to any ISI also clearly increases. Similar changes were observed when sorting based on the first ISI (Fig. S1).

**Figure 1.**
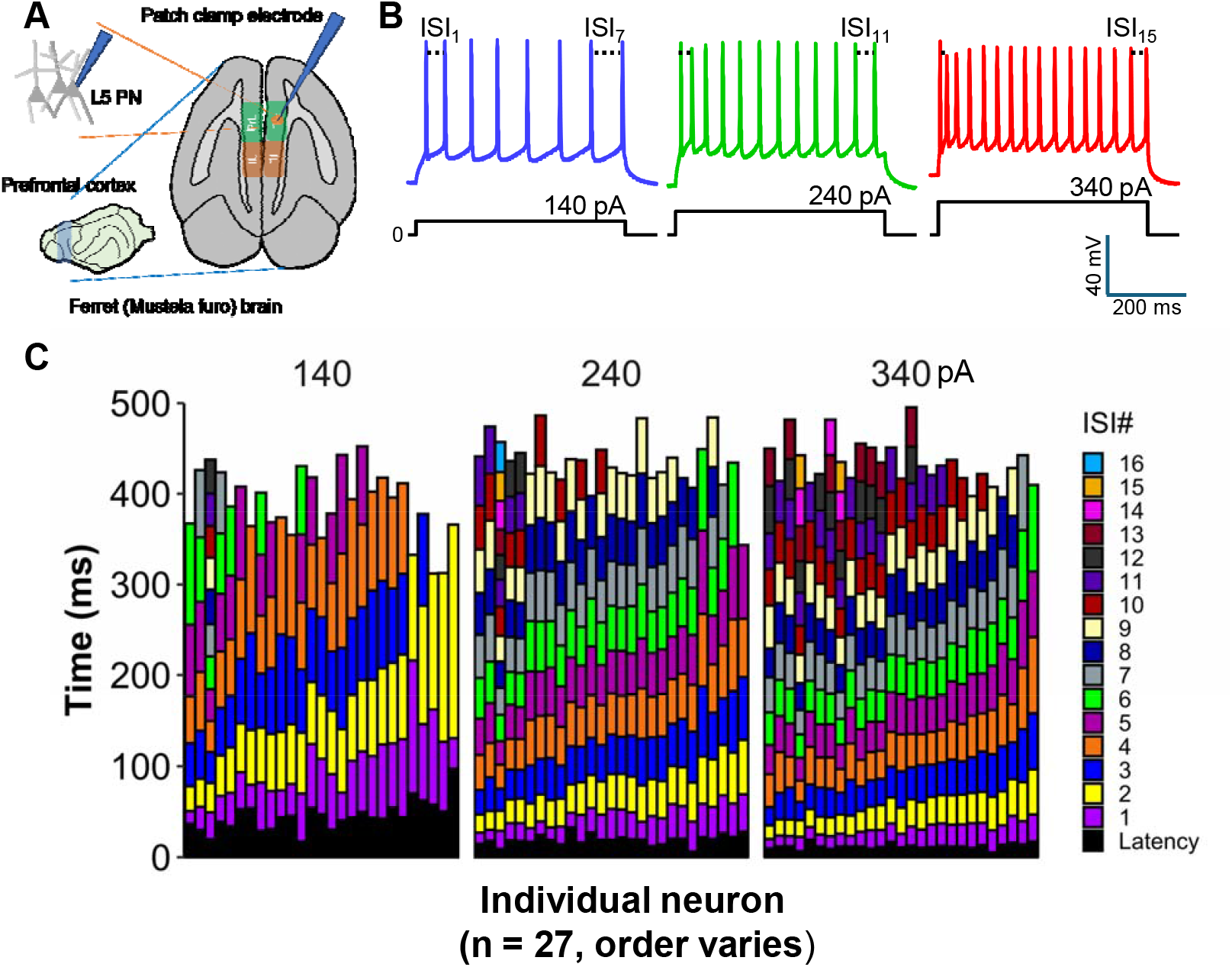
Collection of action potential spiking from layer V pyramidal neurons. *in the medial prefrontal cortex (mPFC)*. **(A)** The patch clamp technique recorded the whole cell potential of layer V pyramidal neurons (PN) in the ferret prelimbic cortex (PrL). **(B)** Representative spike traces evoked by 140 (blue), 240 (green), and 340 (red) pA step current injections for 500 ms were recorded in the same neuron. Black dash lines indicate interspike interval (ISI). **(C)** The cumulative ISI of individual neurons at corresponding stimulation is shown starting from latency (bottom, black) and ISI_1_ (purple) to ISI_16_ (light blue). Cells (*n = 27*) are arranged by their 2^nd^ ISI at the indicated stimulus (see the explanation in the results section), and the order varies for each stimulus.

To confirm this, we calculated the Pearson’s correlation coefficients between the ISI_2_ and the cumulative value up to the fifth ISI for each injected current in as many neurons as possible (Fig. 2A upper; note that some neurons fired fewer than 6 spikes with weaker stimulation). As expected, there was a high correlation between the two for all injected currents. Likewise, all pairs between ISI_1_, ISI_3_, ISI_4_, ISI_5_, and cumulative ISI_5_ (Cum. ISI_5_) showed high Pearson’s correlation coefficients (Fig. 2A, lower). Despite the increase in ISI number the slope of the linear regressions remained similar, indicating that there might be a correlation between each ISI. With the assistance of return plots, we confirmed the strong correlations between the preceding (x-axis) and following (y-axis) ISIs (1st through 6th, Fig. 2B).

**Figure 2.**
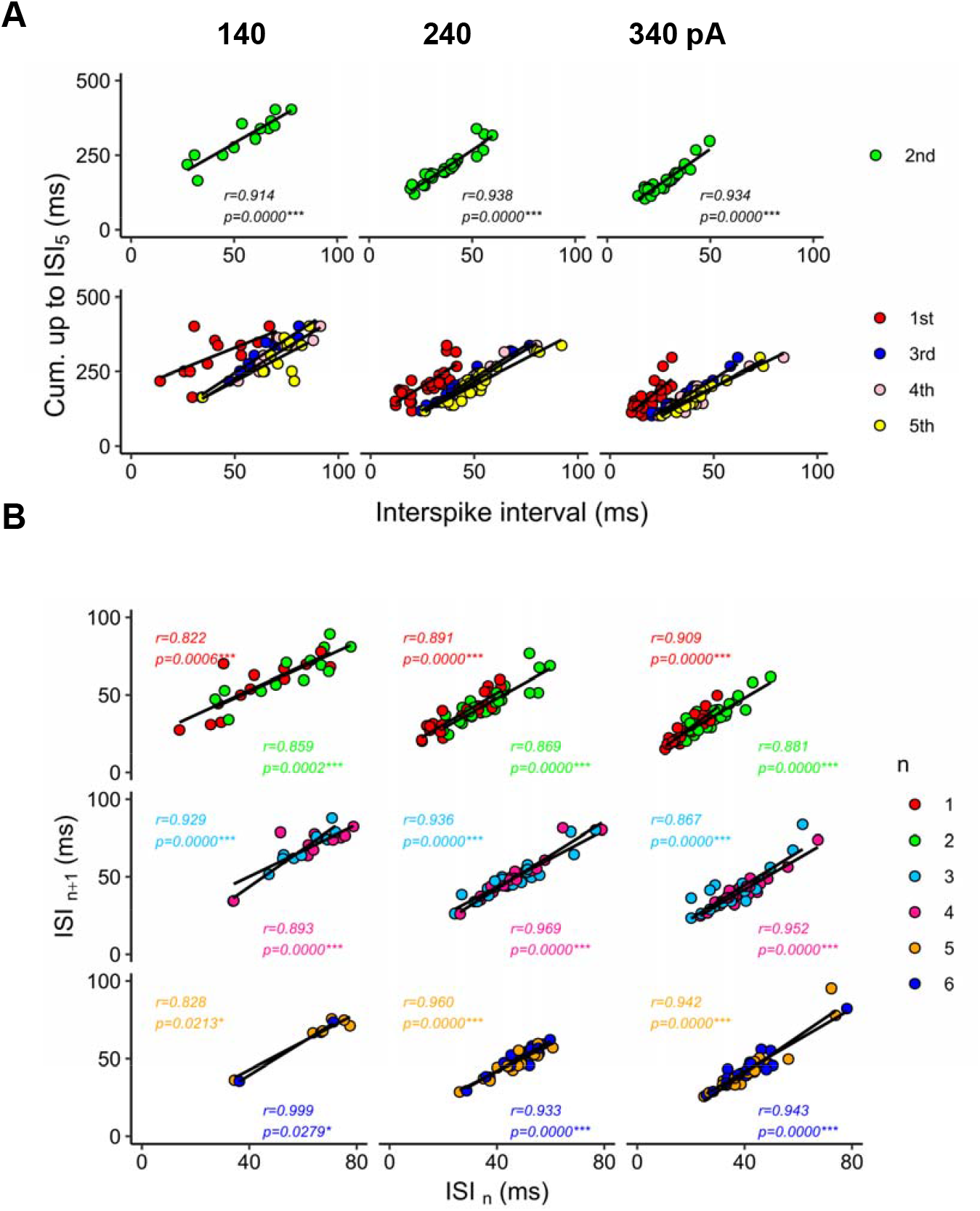
Recurrent correlation between ISIs at selected injected currents (140 pA: left; 240 pA: middle; and 340 pA: right). **(A)** Correlation between ISIs and accumulation up to 5^th^ ISI (ISI_1_+ISI_2_+ISI_3_+ISI_4_+ISI_5_). The ISI_2_ is representatively presented in the upper panels. **(B)** Returning plots showing correlation between ISI_n_ and ISI_n+1_ (n= 1 to 6). Annotations present Pearson’s correlation coefficient *(r)* along with their corresponding *p-values (*p<0*.*05, **p<0*.*01, ***p<0*.*001)*.

Based on these series of strong correlations, this relationship can be expressed as follows:

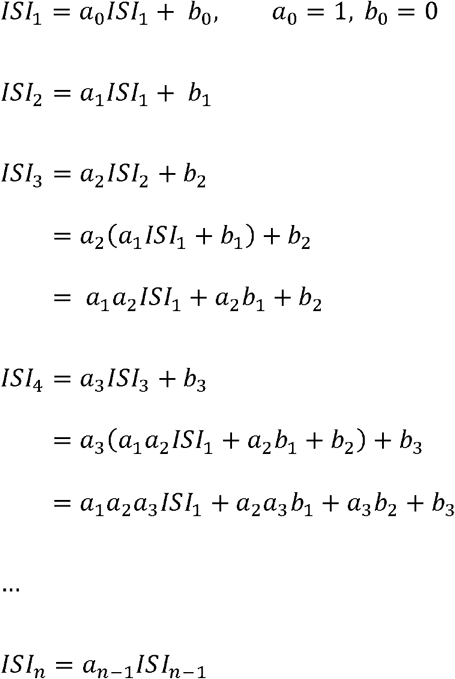

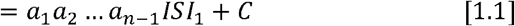

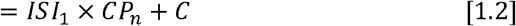

In the above formula, *a*_*n*_ and *b*_*n*_ are the slopes and intercepts of the linear regression line for the preceding and following ISIs (*ISI*_*n*_ *and ISI*_n+1_, respectively), and CP_n_ is the finite product of a_1_ through a_n-1_. This formula consists of a main effect term which is the product of ISI_1_ and the finite cumulative product of slopes, along with a constant term that is composed of combinations of slopes and intercepts. We further examined only the main effect term of [1.1] and [1.2] since the intercepts from linear regression of all combinations of ISI numbers (n) and injected currents (I_inj_) were mainly distributed between −10 and 10 mV, which is relatively smaller than the total ISI value as seen in Fig. S2B. However, the constant term should be studied in the future as its contribution becomes larger as the spike number increases.

Based on *Formula 1*, the cumulation up to the n-th ISI can be expressed as follows:

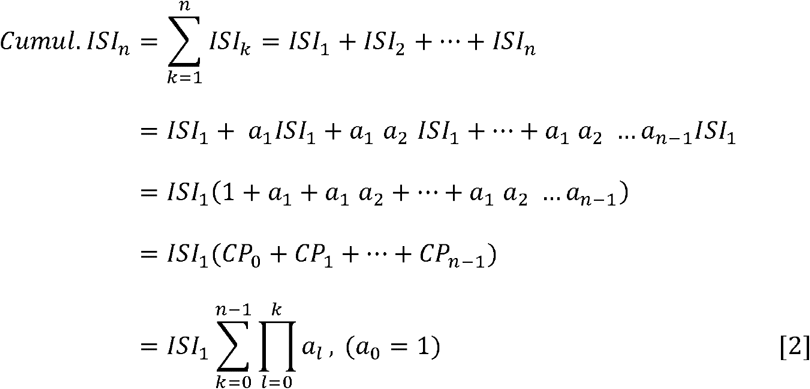

*Formulas 1*.*1, 1*.*2* and *2* are consistent with our observation that there is a correlation between ISIs and their cumulative values.

We investigated how each component of these formulas contributes to constructing the actual ISI in individual neurons. According to Formula 1.2, ISI_1_ is a common coefficient for all following ISIs. In all neurons in the dataset, ISI_1_ was the longest from 21.8 ms to 301.9 ms when the first two spikes were generated at 40-140 pA and was shortest from 10.4 ms to 29.8 ms at 340 pA, the strongest current in our experiment (Fig. 3A). Traces were well-fitted to the negative logarithmic function (Fig. 3B). When *a*_*n*_ (=*ISI*_*n+1*_*/ISI*_*n*_, Fig. 4A upper) and their cumulative product *CP*_*n*_ (Fig. 4A lower) were calculated for individual neurons, more interesting results were observed in the sequence segment. Since *a*_*1*_, *a*_*2*_, *a*_*3*_, and *a*_*4*_ were much larger than 1 across all neurons, they contributed greatly to increasing *CP*_*n*_. However, when n > 4, *a*_*n*_ mostly converged to a number close to 1, there were no large changes in all current intensity levels. Interestingly, the time duration between peak and fast afterhyperpolarization potential (peak-fAHP duration) of preceding spike showed the same trend as *CP*_*n*_ for ISI order (Fig. 4B upper, Fig. S3A and B), but the *fAHP* amplitude did not (Fig. S4). Note that *CP*_*n*_ is equivalent to the normalized ISI (Fig 4B, lower) according to the derived formula. Therefore, the degree of SFA might be related to spike order rather than time after stimulation, regardless of current intensity. Even as the injection current increased, *a*_*n*_ and *CP*_*n*_ at each order (n = 1, 2, 3, 4) barely changed for each neuron, particularly in the range over 200 pA (Fig. 4C). An anticipated mechanism will be discussed later in this paper.

**Figure 3.**
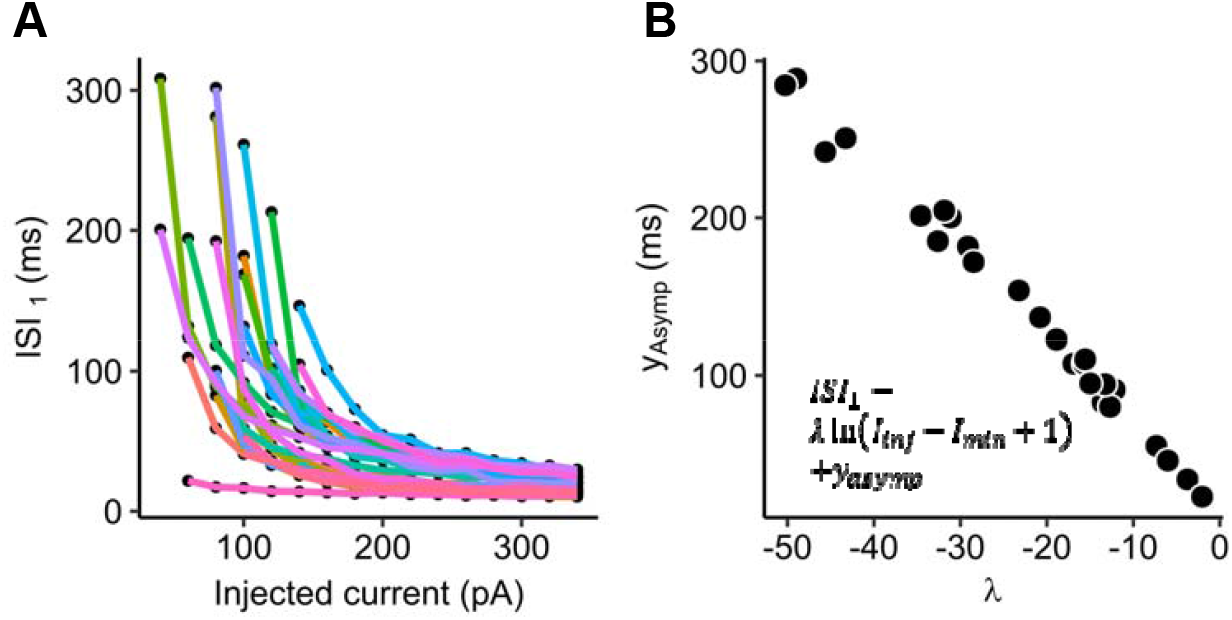
Characteristics of scale factor. **(A)** Change of the 1^st^ ISI upon alteration of the injected current. Dots linked with a line indicate the change in an individual neuron. **(B)** Coefficient of log term and ISI asymptote of curve fittings fitted to indicated formula.

**Figure 4.**
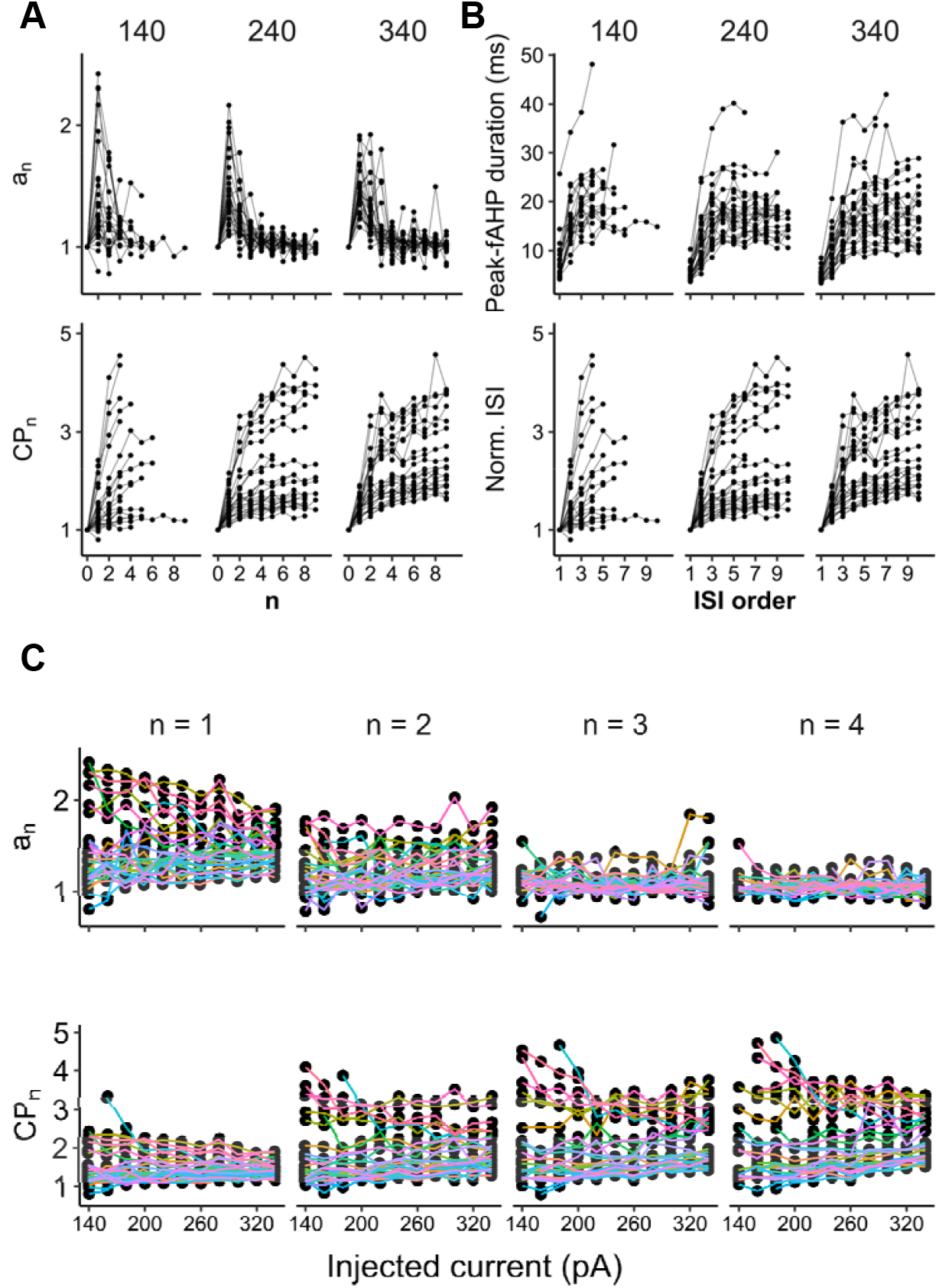
Characteristics of the mathematical sequence. Dots linked with a line indicate the change in an individual neuron. **(A)** Change in a_n_ (upper left), cumulative product of coefficients (lower left), peak-to-fAHP duration (upper right), and normalized ISI (ISI_m_/ISI_1_, lower right) as the number of n (0 to 9) or ISI order (1 to 10) increases at selected stimuli. **(B)** Coefficient (a_1_ to a_4_) and cumulative product of coefficients (CP_1_ to CP_4_) versus current stimuli (140 pA – 340 pA).

Hence, in a spike train evoked by a constant current injection, an ISI between any two spikes can be determined by a timing precursor that exhibits discrete characteristics and a scale factor that varies continuously with the input intensity. Therefore, *Formula 1* can be denoted as below:

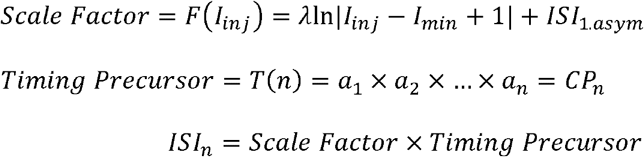

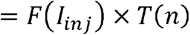

Where *I*_*inj*_ is injected current, *I*_*min*_ is the minimum current that can generate two spikes, and *ISI*_*1,asym*_ is the asymptotic value of ISI_1_, herein ISI_1_ at maximum current input. *n* is spike order represented as an integer greater than 0.

Since the timing precursor rarely changed in a neuron even with different current injections, we were able to simulate the spike train generated by a strong stimulus with the partial information obtained from spike trains generated by weak stimuli. We synthesized 5th through 10th *CP*_*n*_ based on 1st through 4th *CP*_*n*_ extracted from the spike train generated at weaker current injections (200 pA – 300 pA) in individual neurons. *CP*_*5*_ through *CP*_*10*_ were calculated as below:

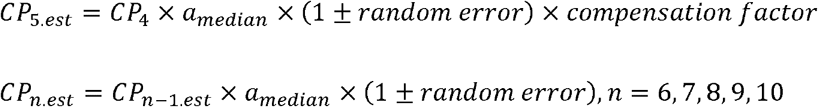

The subscript *est* stands for estimation and *a*_*median*_ stands for median of *a*_*n*_ (5 ≤ n ≤10) of a neuron. The median value of *a*_*n*_ was near 1 across these ISI numbers (Fig. 5A) and ±0.005 (0.5%) of random error was applied to all steps (Heeger, 2000). Also, a compensation of 0.045 (4.5%) was empirically applied to the estimation of *CP*_*5*_ so that the center of the distribution of the entire residual became 0 (Fig. 5B and C). These *CP*_*n*_ were multiplied by the scale factor at 340 pA to reconstruct the spikes and were then compared to the actual spike timing at 340 pA. Figure 5D shows extracted and synthesized timing precursors (*CP*_*n*_) in a sample neuron. We observed that both measured (thick lines) and predicted (thin lines) timing precursors from weak stimuli closely matched those from 340 pA (thick black line). Figure 5E demonstrates that spike trains from a neuron have common components regardless of the input intensity. The timing precursors from weaker stimuli were uniformly multiplied by 24.8 ms, the scale factor (*ISI*_*1*_) of this neuron at 340 pA, and spikes were displayed on the raster plot. In both the measured part (blue rectangle) and the predicted part (green rectangle), most reconstructed spikes from lower stimuli closely followed the actual spikes. On the contrary, when predictions were made from *CP*_*1*_ to *CP*_*3*_ or *CP*_*1*_ to *CP*_*5*_, the former underestimated the actual values, whereas the latter yielded comparable results.

Next, we investigated the theoretical relevance of the two-factor multiplication approach. The open probability (P_O_) of ion channels modulating spike timing can increase cumulatively as spikes repeat, through Ca^2+^ -dependent or -independent ways. Those provide more ion current to the initial leak current until the next spike fires. Larger potassium (or chloride) leak current between spikes will result in a longer time to raise the membrane potential. To confirm this, we assessed the change in spike timing over the increase of the ion conductance using a passive model with HH-like conductance, as described briefly below:

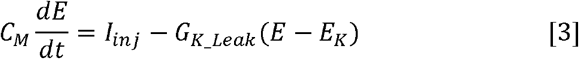

Where *C*_*M*_ (200 pF) is the cell capacitance, *I*_*inj*_ is the injected current, and *G*_*K_Leak*_ and *E*_*K*_ are the potassium leak conductance and reversal potential (−77 mV), respectively. To find the relationship between the change in G_K_ and time required for appropriate membrane potential increase, the following approach was taken. First, the time-voltage change was obtained from Formula 3 using the Runge-Kutta (RK4) method. Next, the time required to produce a specific voltage change with an increase in G_K_ was obtained using linear interpolation. The calculations showed that the time needed to achieve the same voltage change is linearly proportional to the percent change of G_K_ across all given voltage changes (Fig. 6). This might be the basis for the timing precursor and scale factor concept. From Formula 3, the relationship between the injected current (I_0_) and time between spikes can be derived.

**Figure 5.**
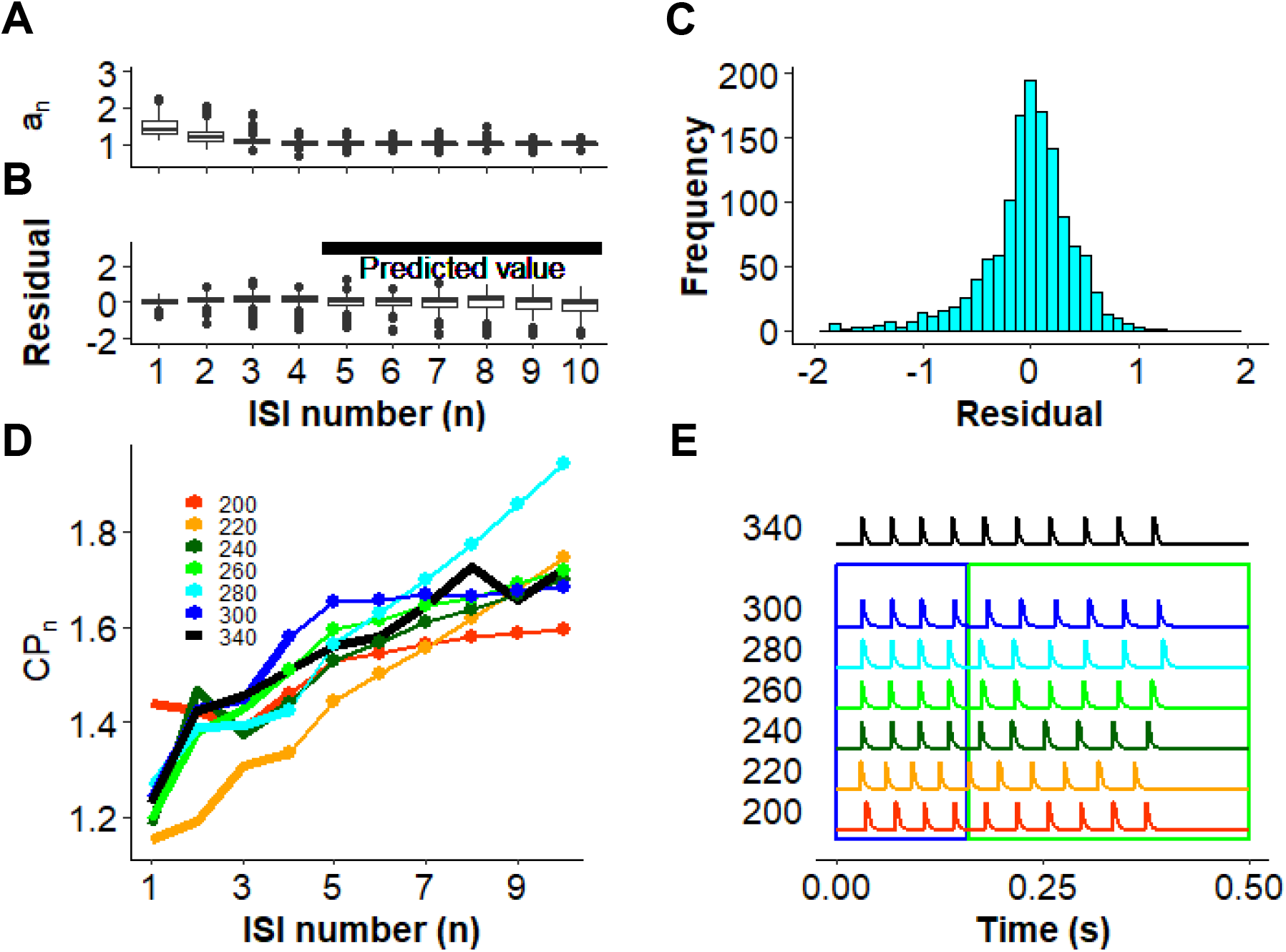
Mimicking fast spike trains with spike trains generated by weaker current stimuli. **(A)** Overall median value of observed *a*_*1*_ through *a*_*4*_ and predicted *a*_*5*_ through *a*_*10*_ of all spikes. Box indicates 1^st^ interquartile, median, and 3^rd^ interquartile. Whiskers indicate 1.5. **(B)** Average residuals of *CP*_*5*_-*CP*_*10*_ against those of 340 pA. **(C)** Distribution of residual of *CP*_*n*_ for n = 5 through 10. **(D-E)** Demonstration of the model in an example neuron. **(D)** *CP*_*5*_ through *CP*_*10*_ (thin line) were predicted from *CP*_*1*_ through *CP*_*4*_ (thick line) of spike trains generated by 200 pA to 300 pA. For the 340 pA trace (thick black line), actual *CP*_*n*_ were plotted for comparison. **(E)** Action potential spike timing was reconstructed by multiplying each term of *CP*_*n*_ obtained in **(D)** by ISI_1_ of 340 pA. 4.5% of compensation was applied to *CP*_*5*_ calculation and 0.5% of random error was applied to every step. Spikes in the blue rectangle indicate reconstruction based on measurement, while spikes in the green rectangle indicate reconstruction based on prediction.

**Figure 6.**
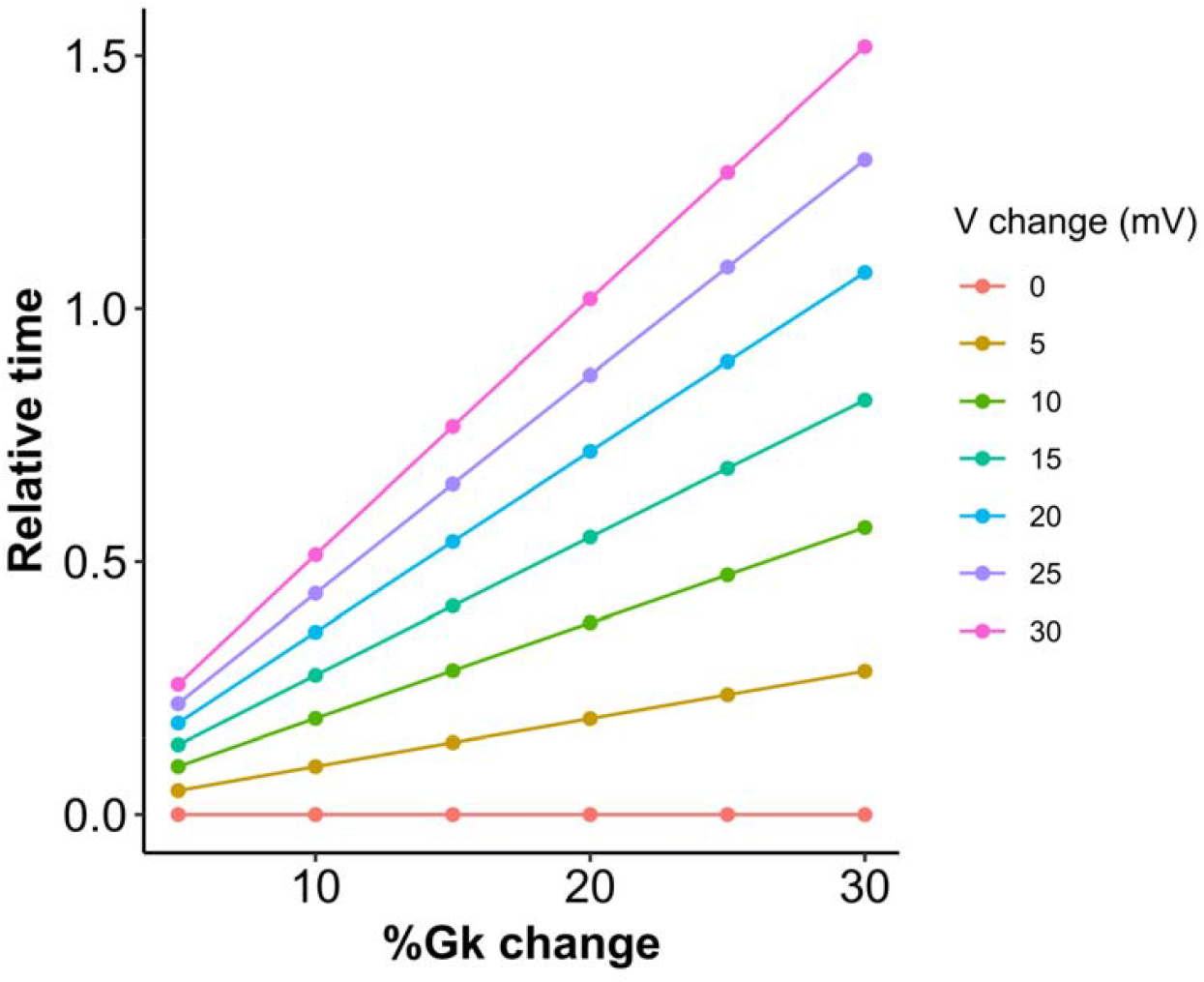
Time for voltage changes regarding % change of potassium conductance.

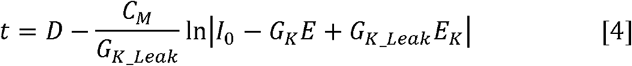

Formula 4 was consistent with our observation that current intensity and ISI have a negative logarithmic relationship.

From Formula [3], the net potassium leak current can be considered as a linear combination of the individual elemental factors. The activation of type-1 muscarinic acetylcholine receptor (mAChR1 or M_1_R) induces the hydrolysis of phosphatidylinositol 4,5-bisphosphate (PIP_2_) through the activation of phospholipase C (PLC), resulting in the release of Ca^2+^ from intracellular storage. It appears to affect spike timing through KCNQ (Kv7) channels regulated by PIP_2_ and calcium-activated potassium channels also regulated by the both Ca^2+^ and PIP_2_ (Suh and Hille, 2008), forming a complex system. We simulated the effect of muscarinic stimulation on spike timing (Fig. 7). The net potassium leak current can be expressed as shown below.

**Figure 7.**
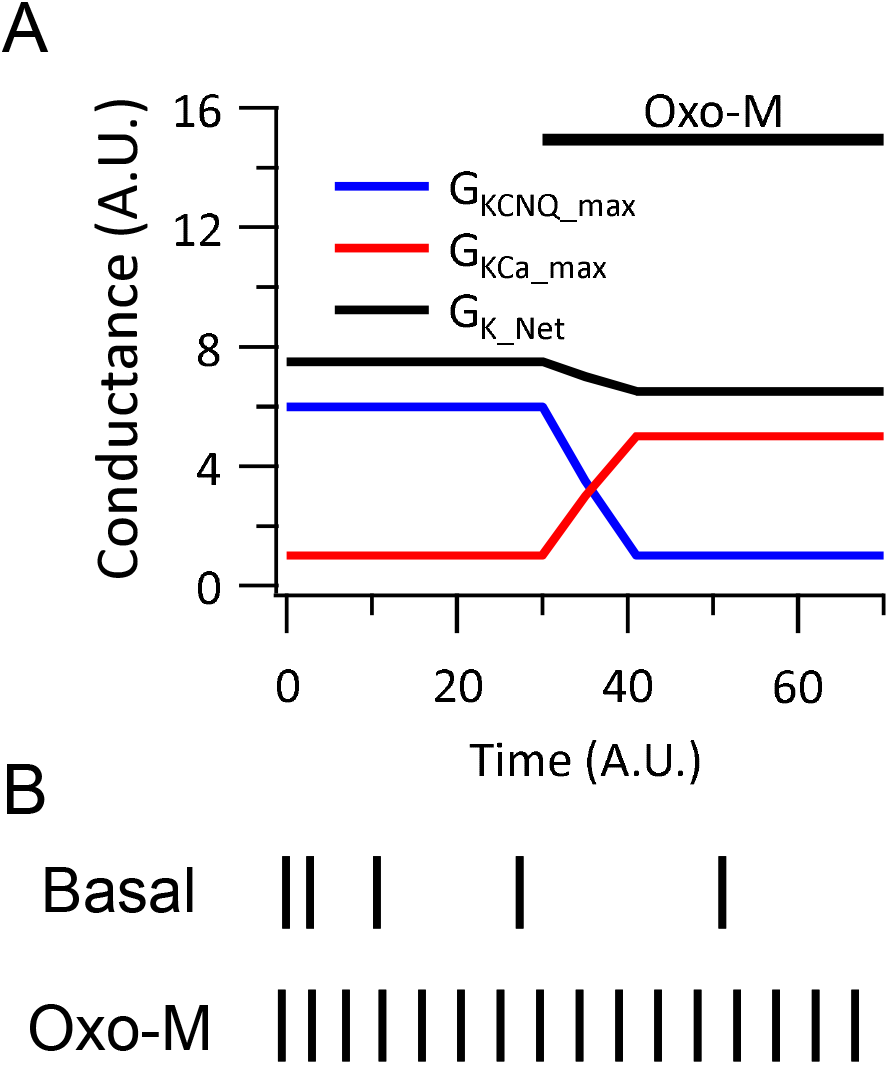
Simulated change of spike patten by the muscarinic stimulation. **(A)** Net G_K_ change (black line) induced by the muscarinic stimulation in PN neurons 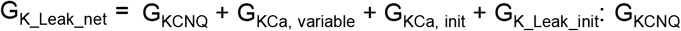 cumulatively increased from 0 until G_KCNQ_max_ was reached. On the other hand, G_KCa_ has a fixed portion affected by cytosolic Ca^2+^ and a variable portion affected by Ca^2+^ influx. **(B)** Raster plots showing presumed spike timing in response to a constant current injection. Spike timings were generated using a ‘sequence model’ reflecting G_KCNQ_ and G_KCa_ values presented in **A**. The effects of the cell capacitance (C_Cell_), the scale factor, was not considered. Calculations can be found in **Table S1**.

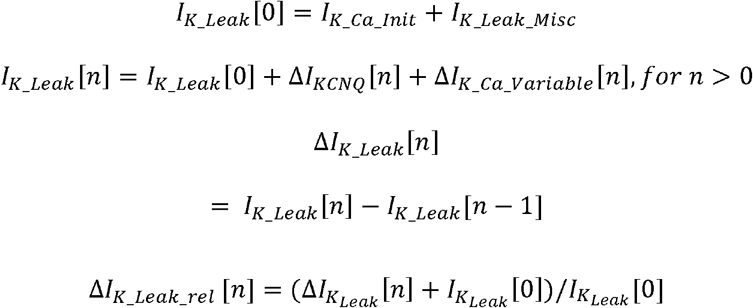

Where *I* _*K_Ca_Init*_ and *I* _*K_L ak_Misc*_ are the basal potassium leak current through calcium dependent potassium channels and miscellaneous potassium channels, respectively. From the linear relationship, *a*_*n*_ and *CP*_*n*_ can be described as below:

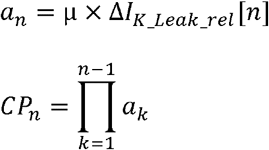

Where is a proportional constant between *a*_*n*_ and 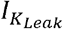 Assumptions are that 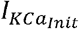 relies on the initial calcium level at each condition, and 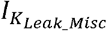 was fixed at 0.5 *I*_*KCNQ*_ and *I ca_variable* cumulatively increase through 1^st^ to 4^th^ spike until they reach their maximum conductance. The net leak current decreased after the muscarinic stimulation, considering the PIP_2_-channel interaction on *I* _*KCa*_ (Zhang et al., 2014; Harraz et al., 2020). We put the parameters into the sequence model (Table 1 and 2). In the basal condition, there was a stronger SFA due to the larger portion of *I*_*KCNQ*_ to *I*_*K_Leak*_. However, larger *I*_*KCa*_ induced by the increased cytosolic calcium results in shorter and rarely changed spike pattern upon the activation of M_1_R. This result was comparable to the actual experimental results conducted in other laboratories (Carver and Shapiro, 2019). Additionally, this phenomenon was observed in current clamp recordings of PN neurons in mouse V1 and ferret and mouse mPFC, suggesting that it may also occur in PN neurons in other regions.

**Table 1.** Datapoints for basal condition used in Figure 6B. The relative spike time was achieved by stepwise manipulation of each element. I_K_Leak_ = I_KCa_Init_ +I_K_Leak_Misc_ + I_KCNQ_ + I_KCa_Variable_. I_KCa_Init_ and I_K_Leak_Misc_ represent the initial, and fixed leak current formed by calcium-activated potassium channels and miscellaneous potassium channels, respectively. I_KCNQ_ is potassium current formed by KCNQ channel. I_KCa_variable_ is the volatile calcium-activated potassium current increased by the influx of external calcium. Relative change was obtained by adding I_K_Leak_[0] to ΔI_K_Leak_. a_n_ is obtained by applying a proportional constant 1 to I_K_Leak_, a_n_normal_ is a_n_/a_0_, and CP_n_ is a finite product of a_n_normal_. Spike time is a cumulation of CP_n_, and Reletive_spike_time is achieved by multiplying a_0_ to the spike time.

| Spike# | $I_{KCa\_Init}$ | $I_{KCa\_var}$ | $\Delta I_{KCa\_variable}$ | $I_{K\_Leak\_Misc}$ | $I_{KCNQ}$ | $\Delta I_{KCNQ}$ | $I_{K\_Leak}$ | $\Delta I_{K\_Leak}$ | Relative_Change | $a_n$ | $a_{n\_normal}$ | $CP_n$ | SpikeTime | Reletive_spike_time |
| --- | --- | --- | --- | --- | --- | --- | --- | --- | --- | --- | --- | --- | --- | --- |
| 0 | 0.5 | 0 | 0 | 0.5 | 0 | 0 | 1 | 0 | 1 | 1 | 1 | 1 | 0 | 0 |
| 1 | 0.5 | 0.125 | 0.125 | 0.5 | 3 | 3 | 4.125 | 3.125 | 4.125 | 4.125 | 4.125 | 4.125 | 4.125 | 4.125 |
| 2 | 0.5 | 0.25 | 0.125 | 0.5 | 4.7 | 1.7 | 5.95 | 1.825 | 2.825 | 2.825 | 2.825 | 11.65313 | 15.778125 | 15.778125 |
| 3 | 0.5 | 0.375 | 0.125 | 0.5 | 5.7 | 1 | 7.075 | 1.125 | 2.125 | 2.125 | 2.125 | 24.76289 | 40.54101563 | 40.54101563 |
| 4 | 0.5 | 0.5 | 0.125 | 0.5 | 6 | 0.3 | 7.5 | 0.425 | 1.425 | 1.425 | 1.425 | 35.28712 | 75.82813477 | 75.82813477 |
| 5 | 0.5 | 0.5 | 0 | 0.5 | 6 | 0 | 7.5 | 0 | 1 | 1 | 1 | 35.28712 | 111.1152539 | 111.1152539 |
| 6 | 0.5 | 0.5 | 0 | 0.5 | 6 | 0 | 7.5 | 0 | 1 | 1 | 1 | 35.28712 | 146.402373 | 146.402373 |
| 7 | 0.5 | 0.5 | 0 | 0.5 | 6 | 0 | 7.5 | 0 | 1 | 1 | 1 | 35.28712 | 181.6894922 | 181.6894922 |
| 8 | 0.5 | 0.5 | 0 | 0.5 | 6 | 0 | 7.5 | 0 | 1 | 1 | 1 | 35.28712 | 216.9766113 | 216.9766113 |

**Table 2.**
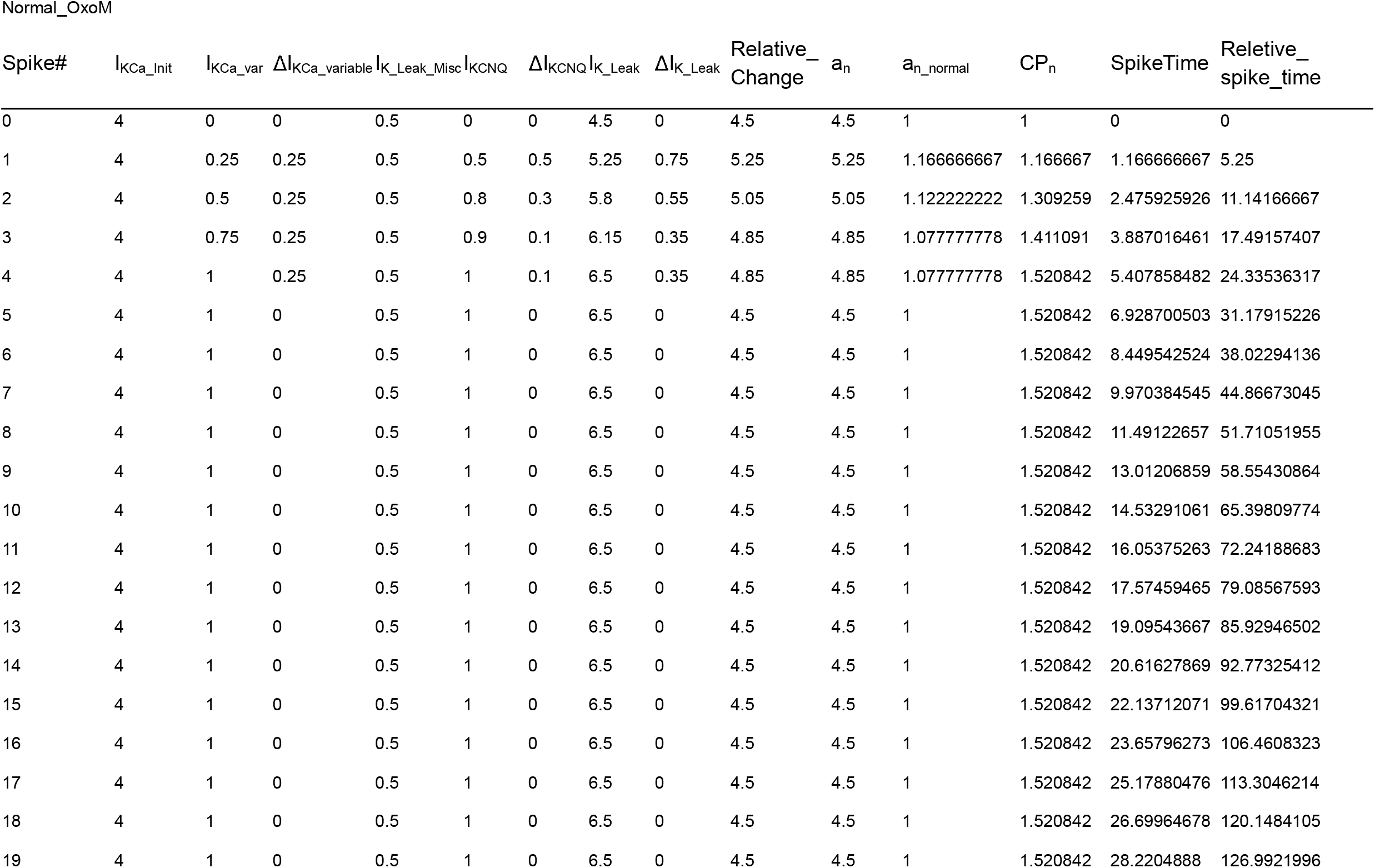
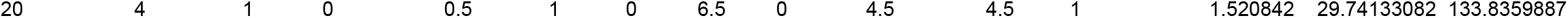
Datapoints for action potential timing after muscarinic stimulation used in Figure 6B. The relative spike time was achieved by multiplying 4.5 (value of a_0_) to Spike Time.

## Discussion

Considering the series of strong correlations observed between preceding and following ISIs, we derived a formulaic relationship, thereby elucidating the two elements contributing to the formation of action potential formation: the sequence component as a timing precursor, and the first ISI as a scale factor. Those two factors originated from ion channel activities and passive membrane properties. Subsequently, we observed a strong regularity in occurrence of tonic spike trains and proposed a model for predicting spike firings by leveraging the characteristics of these two elements, which depend on the spike order and the intensity of stimulation, respectively. We also demonstrated that this approach could reflect the activity of each subfactor to the spike timing, based on their linearity.

The SFA is a non-regularized process that alters the spike interval and adds complexity to spike analysis. SFA is mediated by ion channels such as KCNQ (M-) channels (Gu et al., 2005), Anoctamin 2 (ANO2; Ha et al., 2016; Ha and Cheong, 2017) calcium activated chloride channel, small (SK; Engel et al., 1999) and large (BK; Gu et al., 2007) conductance calcium-activated potassium channels, and hyperpolarization activated hyperpolarization-activated cyclic nucleotide gated (HCN) channels (Gu et al., 2005), which are widely expressed in the brain. Pharmacological inhibition of these channels decreases or abolishes fast and medium after hyperpolarization (AHPs; Gu et al., 2005; Peng et al., 2017), indicating that they contribute to repolarizing conductance (Gu et al., 2005). Repetitive generation of spikes accumulates cytosolic calcium (Oh et al., 2016), causing a unitary increase in the open probability (P_O_) of Ca^2+^-activated K^+^ or Cl^-^ channels (Dayan and Abbott, 2001). It is likely that a similar mechanism, involving the cumulative increase in P_O_ by unitary stimulus, exists in KCNQ channels (Zhang et al., 2022) due to its slow deactivation characteristics (Chen et al., 2015; Hou et al., 2017), but in a Ca^2+^-independent manner. When a constant current stimulus is provided, SFA is prominently observed between early spikes (n≤4, also demonstrated in (Gu et al., 2005)), such that the channel’s open probability increases until saturation is reached (n>4). Therefore, the emergence of the recursive sequence appears to arise from the consistent progression of adaptation across each step, forming a factorial spike history. In another sense, the timing precursor can be considered as an oscillation with initial damping in a discrete step, and SFA is a transient process reaching a steady state.

The ion-channel factors determining spike timing can be added or subtracted due to their linear characteristic. For instance, we did not include the effect of HCN channels or ANO channels in the simulation, but terms for those ion currents (with their appropriate reversal potentials) can be added to the calculation. Thinking the other way around, since the net leak current is expressed as a linear combination of multiple ion channel activities, individual channel activities can be tracked using mathematical methods, e.g., a system of simultaneous equations or series of matrices, with parameters of spike trains generated in different drug conditions.

So far, we have broken down the elements that make up the pattern of tonic spike trains in response to a constant current input and have inferred the origins of each element. With this finding, contributions of both passive membrane properties and ion conductance in the spike patterning are revealed, and this will provide the resolution needed to investigate many neurological disorders arising from abnormal spike dynamics more precisely. Furthermore, our findings can greatly help in understanding the characteristics of neural spiking. Additionally, as neuronal models become more crucial, the need for computational efficiency and management of complexity also rises. The cumulative model we presented is based on the practical phenomenon of SFA in live neurons, making its calculations straightforward and intuitive. Although simpler than previously known models, it may be effective and suitable for application in artificial intelligence through computation (Ganguly et al., 2024), neural signal detection (Ratnam and Nelson, 2000; Nesse et al., 2021), and neural coding research (Prescott and Sejnowski, 2008; Naud and Gerstner, 2012; Lee et al., 2023).

## Acknowledgements

This work was supported by #I01BX005678-01A1 from the United States (U.S.) Department of Veterans Affairs Biomedical Laboratory R&D Service. The contents do not represent the views of the U.S. Department of Veterans Affairs or the United States Government.

We would like to thank Fabrizio Gabbiani and Steve Cox for comments and practical suggestions.

## Author contributions

DK performed patch clamp recordings. DK and KW conceived of the presented idea. DK derived and KW re-examined the formular. DK performed data analysis and wrote most parts of the manuscript with the support of AM. KW wrote part of the introduction and discussion. KW investigated the characteristics of KCNQ channels involved in this study. AM suggested a verification model demonstrated in Figure 5. MP contributed to the final version of the manuscript. All authors discussed the results, provided critical feedback and contributed to the final manuscript.

## Data and code availability

The data that support the finding of this study as well as R and Python codes used for analysis are available at the following URL: https://github.com/ehd12/AP_PATTERN.

