## Supplementary material for "A mathematical sequence representing tonic action potential spike trains": Fig S1, Fig S2, Fig S3, Fig S4

### Slide 1
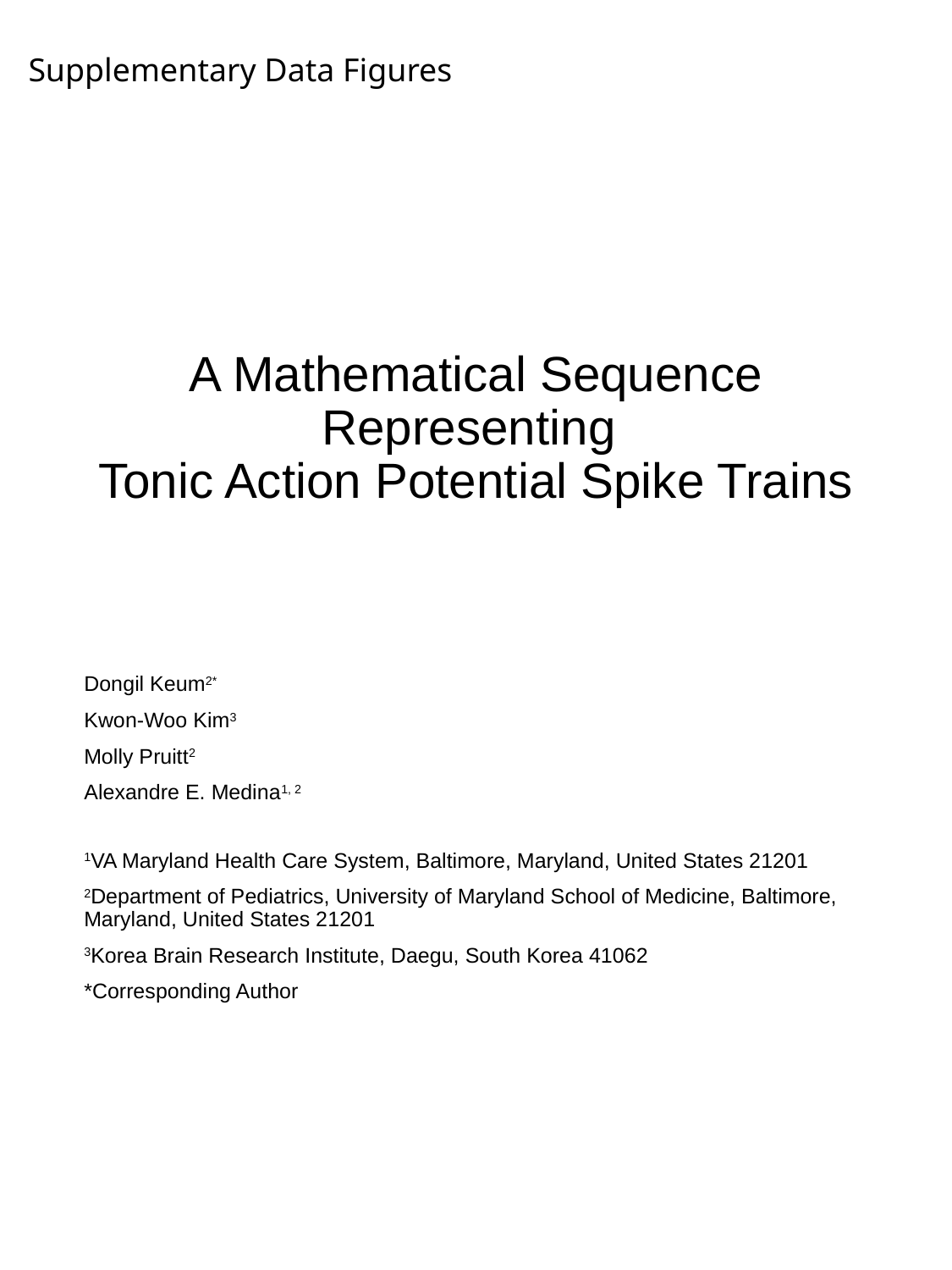

Supplementary Data Figures
#### A Mathematical Sequence Representing Tonic Action Potential Spike Trains
Dongil Keum^2*^
Kwon-Woo Kim^3^
Molly Pruitt^2^
Alexandre E. Medina^1, 2^
^1^VA Maryland Health Care System, Baltimore, Maryland, United States 21201
^2^Department of Pediatrics, University of Maryland School of Medicine, Baltimore, Maryland, United States 21201
^3^Korea Brain Research Institute, Daegu, South Korea 41062
*Corresponding Author

### Slide 2
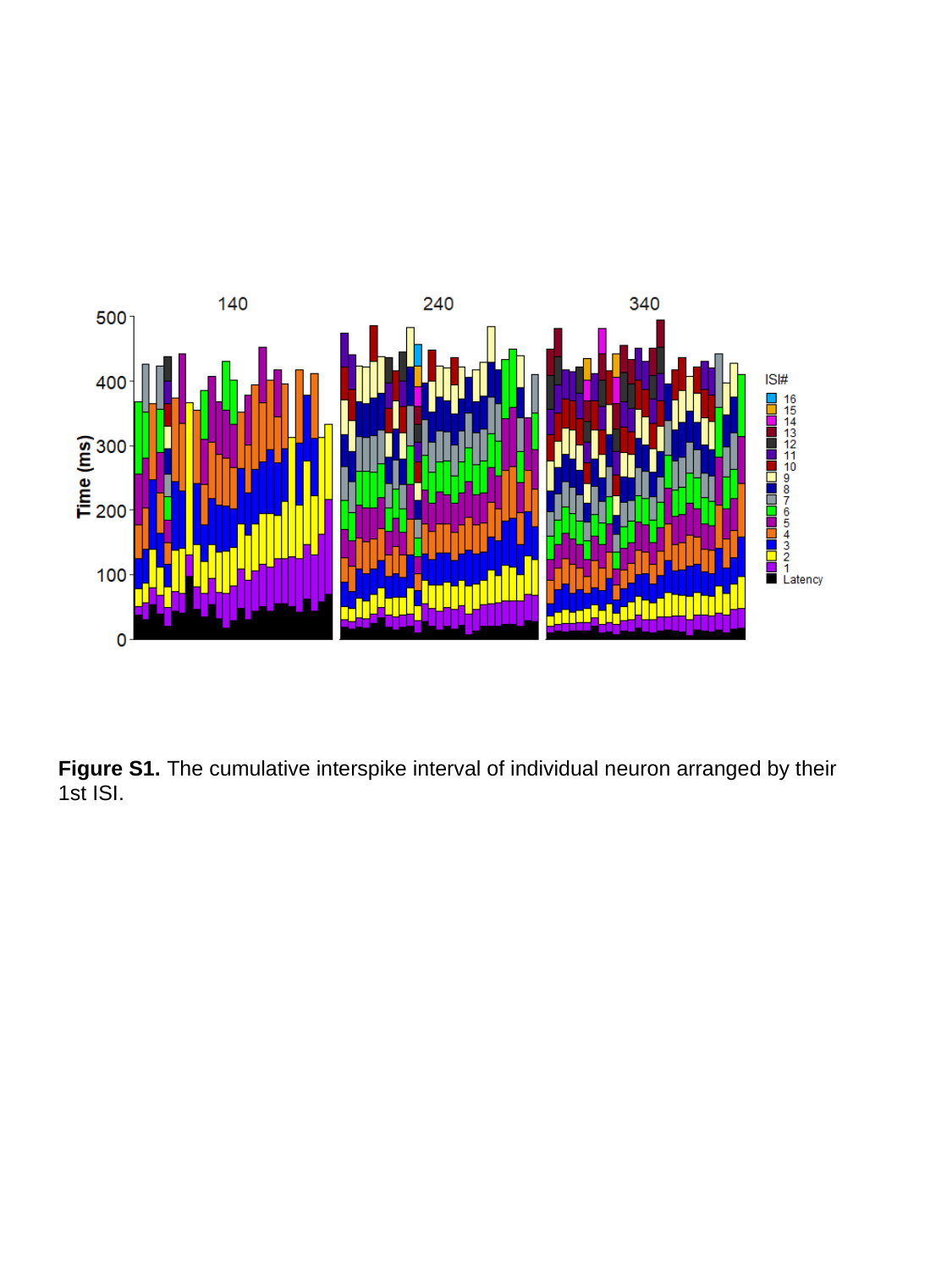

Figure S1. The cumulative interspike interval of individual neuron arranged by their 1st ISI.

### Slide 3
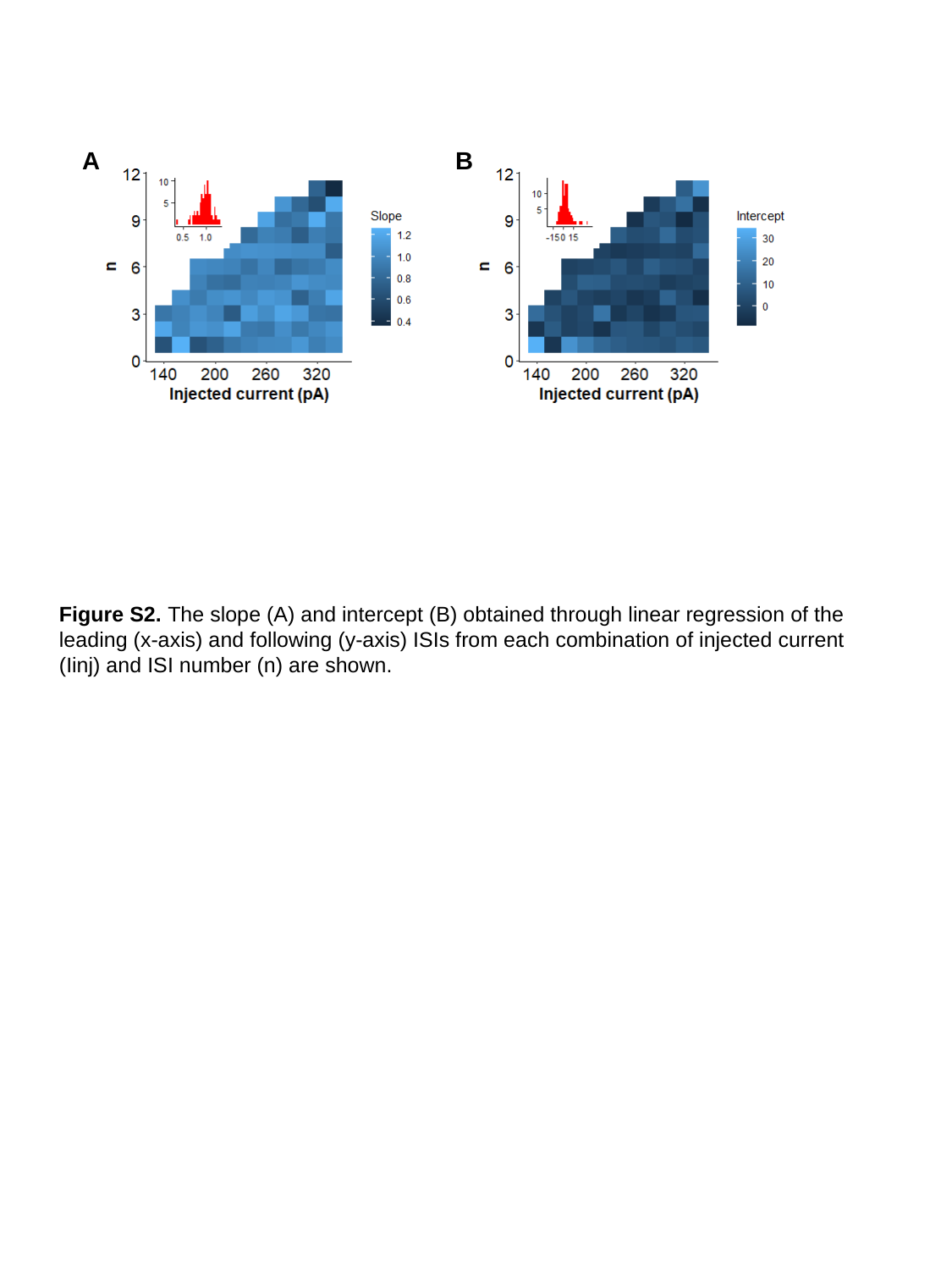

A
B
Figure S2. The slope (A) and intercept (B) obtained through linear regression of the leading (x-axis) and following (y-axis) ISIs from each combination of injected current (Iinj) and ISI number (n) are shown.

### Slide 4
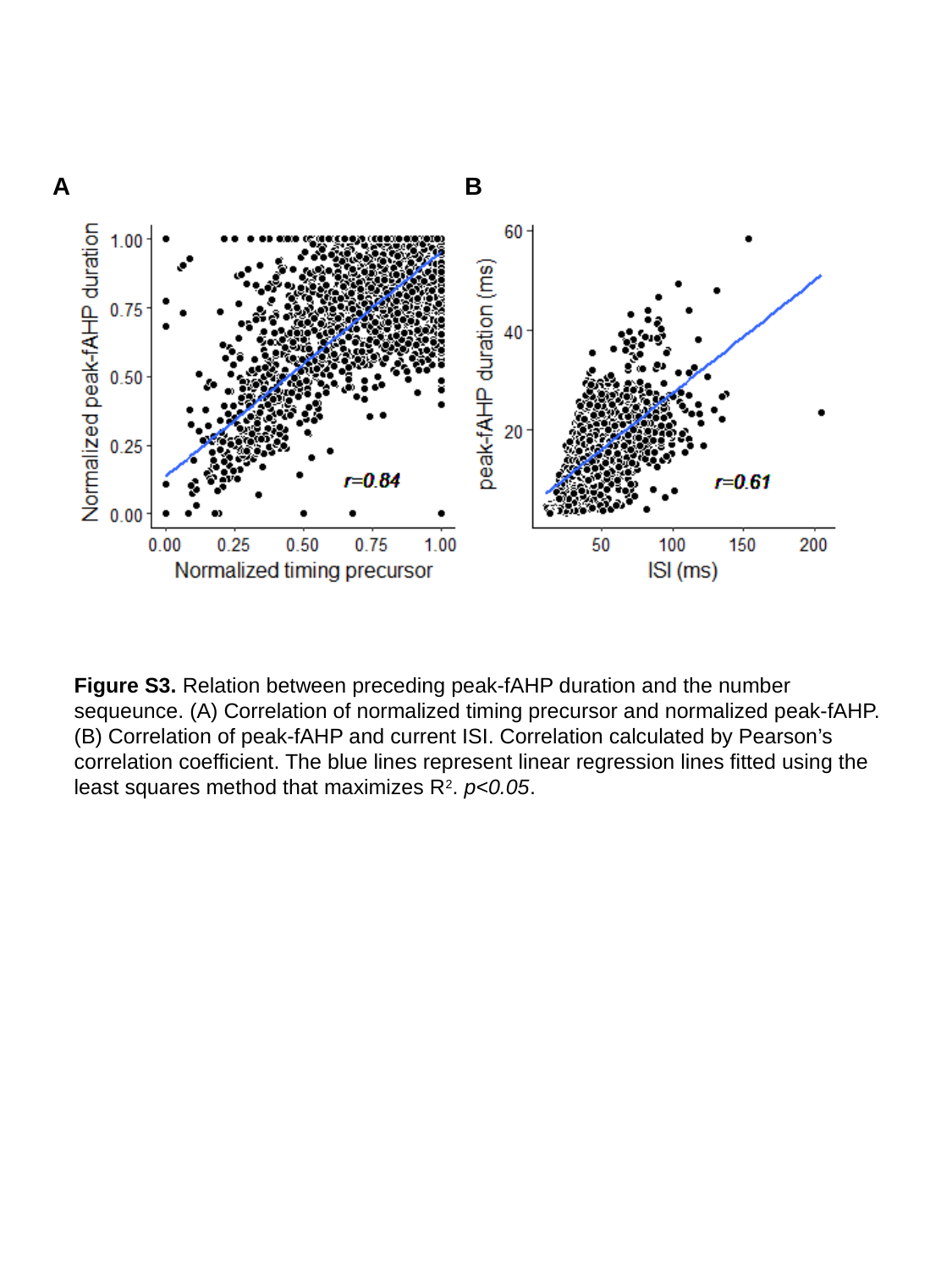

B
A
Figure S3. Relation between preceding peak-fAHP duration and the number sequeunce. (A) Correlation of normalized timing precursor and normalized peak-fAHP. (B) Correlation of peak-fAHP and current ISI. Correlation calculated by Pearson’s correlation coefficient. The blue lines represent linear regression lines fitted using the least squares method that maximizes R^2^. p<0.05.

### Slide 5
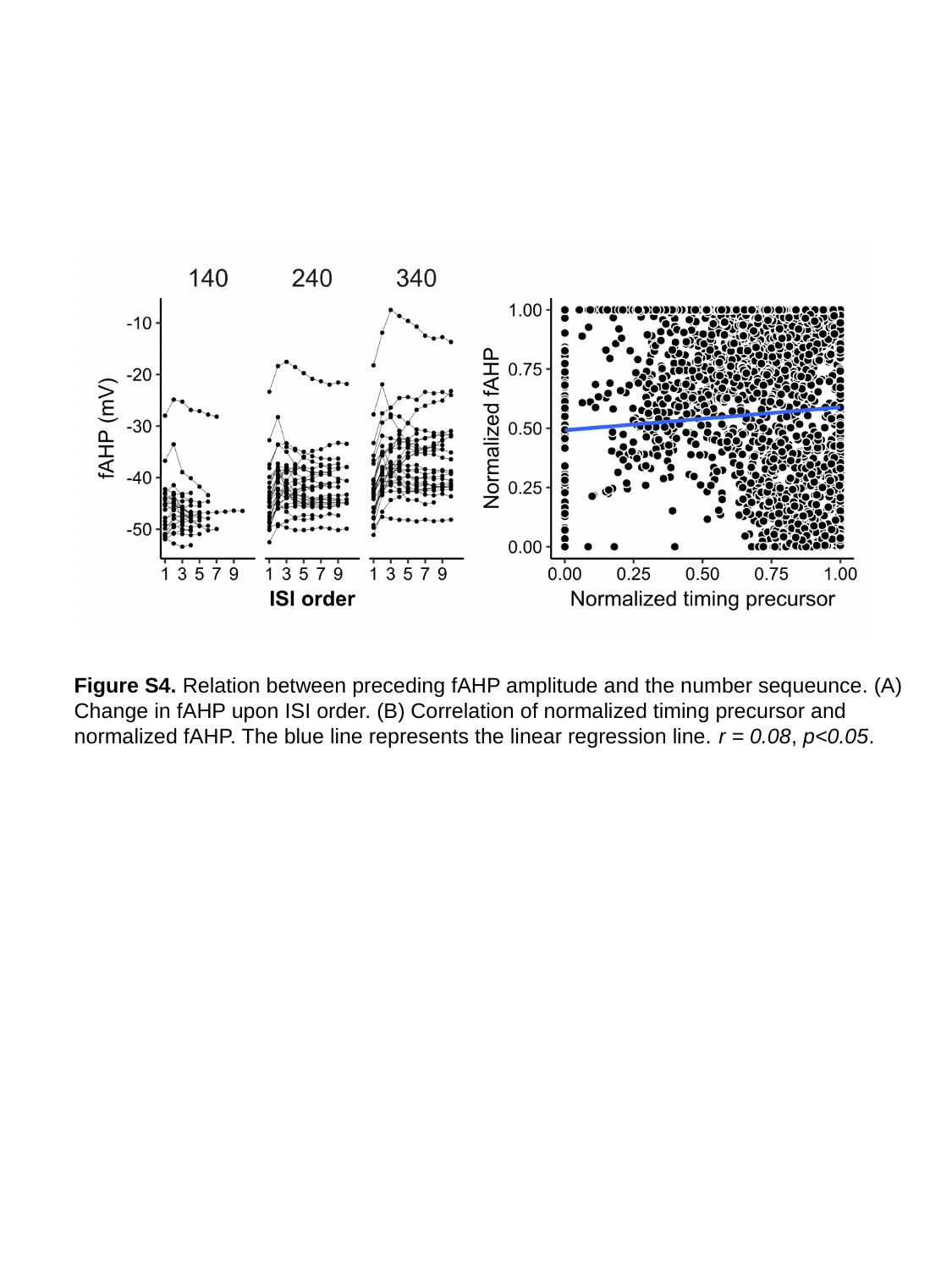

Figure S4. Relation between preceding fAHP amplitude and the number sequeunce. (A) Change in fAHP upon ISI order. (B) Correlation of normalized timing precursor and normalized fAHP. The blue line represents the linear regression line. r = 0.08, p<0.05.
